# Flexible comparison of batch correction methods for single-cell RNA-seq using BatchBench

**DOI:** 10.1101/2020.05.22.111211

**Authors:** Ruben Chazarra-Gil, Stijn van Dongen, Vladimir Yu Kiselev, Martin Hemberg

## Abstract

As the cost of single-cell RNA-seq experiments has decreased, an increasing number of datasets are now available. Combining newly generated and publicly accessible datasets is challenging due to non-biological signals, commonly known as batch effects. Although there are several computational methods available that can remove batch effects, evaluating which method performs best is not straightforward. Here we present BatchBench (https://github.com/cellgeni/batchbench), a modular and flexible pipeline for comparing batch correction methods for single-cell RNA-seq data. We apply BatchBench to eight methods, highlighting their methodological differences and assess their performance and computational requirements through a compendium of well-studied datasets. This systematic comparison guides users in the choice of batch correction tool, and the pipeline makes it easy to evaluate other datasets.

## Introduction

Single-cell RNA sequencing (scRNA-seq) technologies have made it possible to address biological questions that were not accessible using bulk RNA sequencing (1), e.g. identification of rare cell types (2,3), discovery of developmental trajectories (4–6), characterization of the variability in splicing (7–11), investigations into allele specific expression (12–15), and analysis of stochastic gene expression and transcriptional kinetics (11,16). There are currently a plethora of different protocols and experimental platforms available (17,18). Considerable differences exist among scRNA-seq protocols with regards to mRNA capture efficiency, transcript coverage, strand specificity, UMI inclusion, and other potential biases (17,18). It is well known that these and other technical differences can impact the observed expression values, and if not properly accounted for they could be confounded with biological signals (19). Such differences arising due to non-biological factors are commonly known as batch effects.

Fortunately, with appropriate experimental design it is possible to remove a portion of the batch effects computationally, and recently there has been a large degree of interest in developing such methods for scRNA-seq. We group the methods into three categories depending on what space they operate on with respect to the expression matrix (Fig 1a). The expression matrix represents the number of reads found for each cell and gene, and it is central to computational analyses. The first set of methods, mnnCorrect, limma, ComBat, Seurat 3 (hereafter referred to as Seurat) and Scanorama, produce a merged, corrected expression matrix. The second set, Harmony and fastMNN, instead operate on a low-dimensional embedding of the original expression matrices. As such their output cannot be used for downstream analyses which require the expression matrix, limiting their use for some applications. Finally, the BBKNN method operates on the k-nearest neighbor graph constructed from the expression matrices and consequently its output is restricted to downstream analyses where only the cell label can be used.

**Figure 1:**
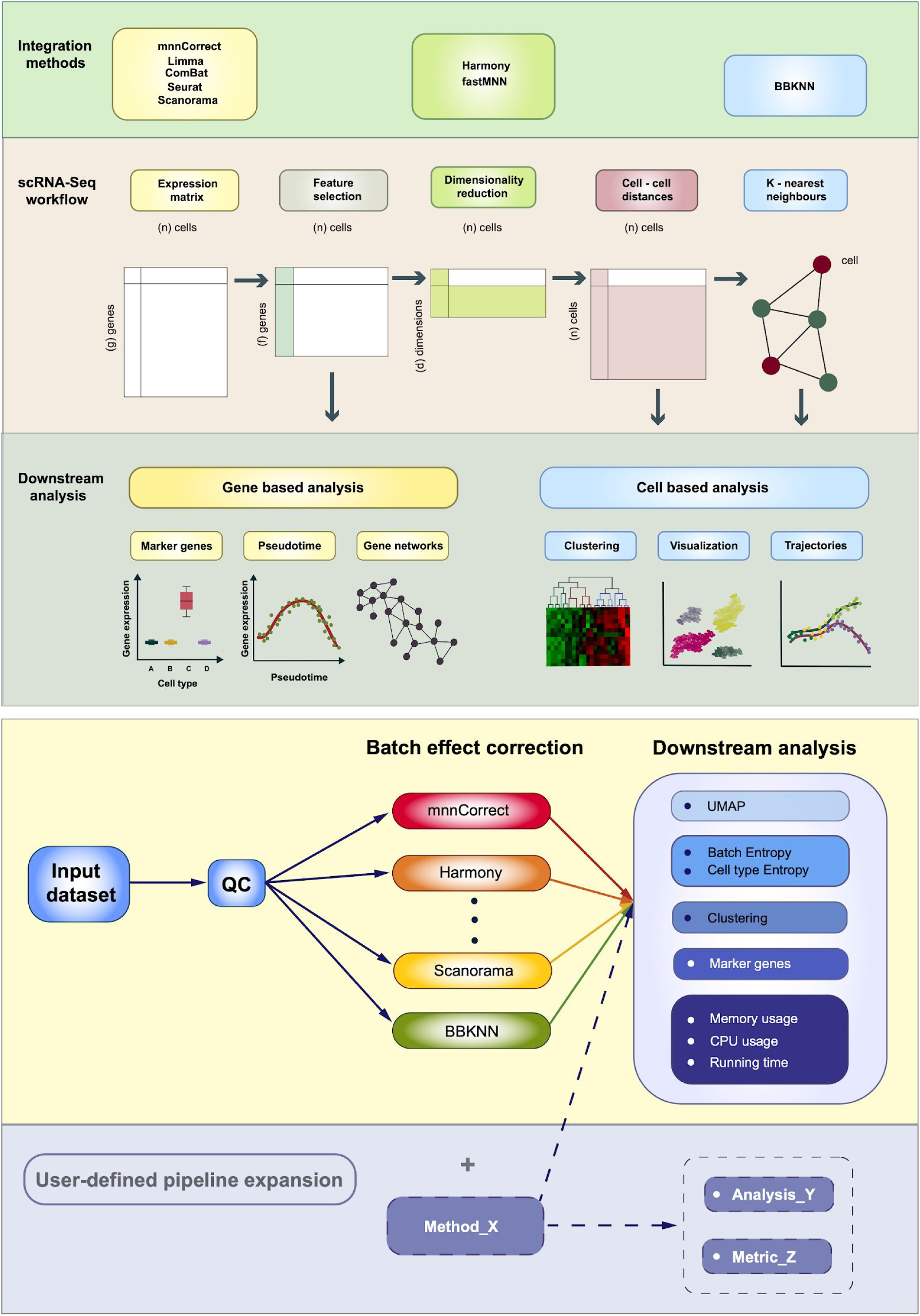
Overview of workflows and batch correction methods. (a) Overview and classification of eight batch effect removal tools. (b) Schematic overview of the BatchBench pipeline.

As the choice of batch correction method may impact the downstream analyses, the decision of which one to use can be consequential. To decide what method to use, most researchers rely on benchmarking studies. Traditionally such comparisons are carried out using a compendium of relevant datasets. The downside of this approach is that methods published after the benchmark was carried out are not included and that the comparison may not have featured datasets that contain all the relevant features required to evaluate the methods. To overcome these issues we have developed BatchBench (Fig 1b), a flexible computational pipeline which makes it easy to compare both new methods and datasets using a variety of criteria. Here we report on the comparison of eight popular batch effect removal methods (Table 1) using three well-studied scRNA-seq datasets. BatchBench is implemented in Nextflow (20) and it is freely available at https://github.com/cellgeni/batchbench under the MIT Licence.

**Table 1:**
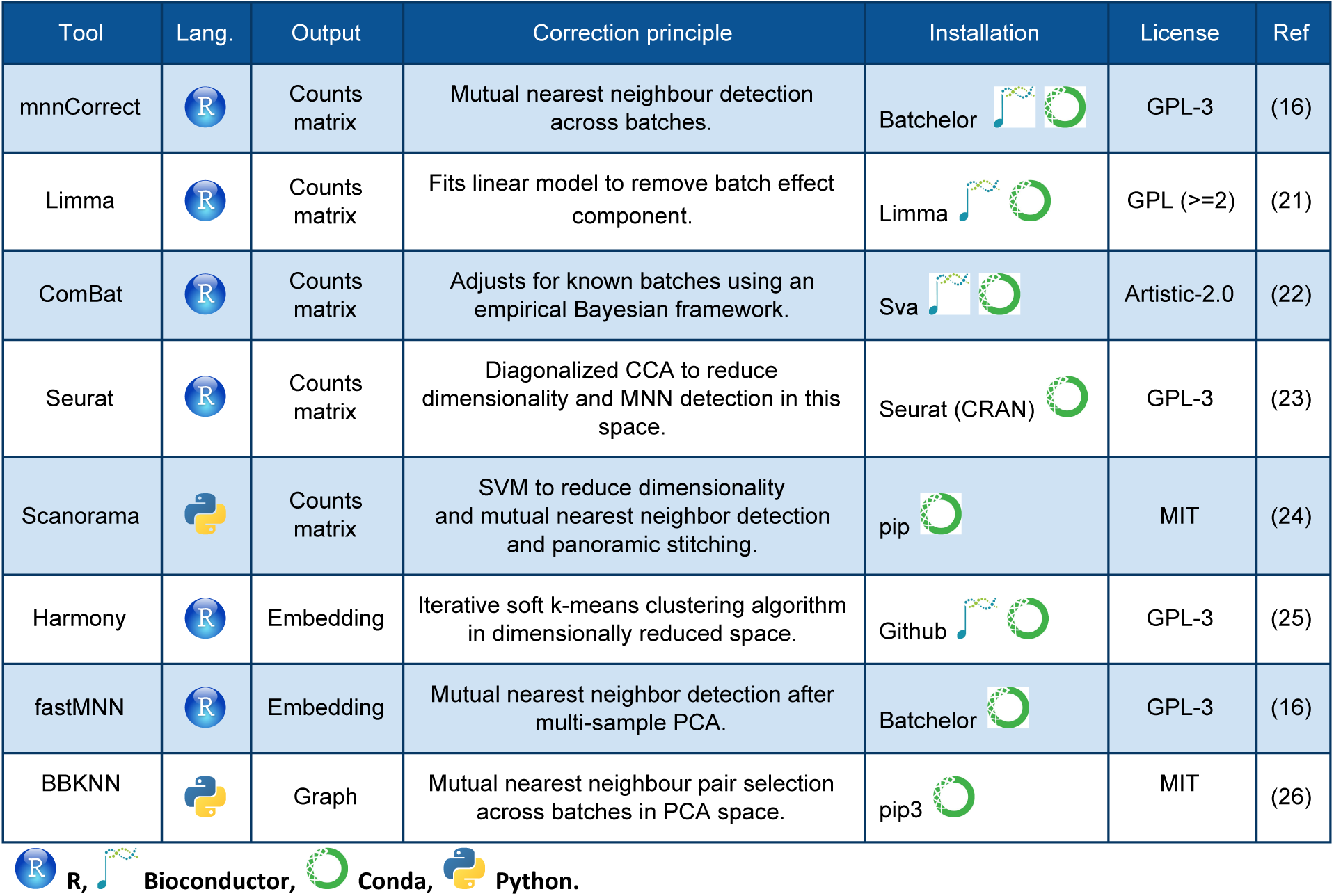
Summary of the eight batch correction methods considered in this study.

By default, BatchBench evaluates batch correction methods based on two different entropy metrics. The normalized Shannon entropy is used to quantify how well batches are aligned while preserving the separation of different cell populations. However, the entropy measures do not provide a complete picture of how the batch correction impacts downstream analyses. Therefore, BatchBench has a modular design to allow users to incorporate additional metrics, and we provide two examples of such metrics - unsupervised clustering and identification of marker genes. Three different unsupervised clustering methods are applied to the merged cells to afford the user a better understanding of how the different methods affect this step which is often central to the analysis. We also compare cell-type specific marker genes to understand how different batch correction methods affect the expression levels.

## Results

### Entropy measures quantify integration of batches and separation of cell types

To illustrate the use of BatchBench we first considered three scRNA-seq studies of the human pancreas (27–29). Even though the samples were collected, processed and annotated independently, several comparisons have shown that batch effects can be overcome (19,30). Visualization of the uncorrected data using UMAP reveals a clear separation of the major cell types across batches (Fig 2a). As expected, all of the methods in our study were able to merge equivalent cell populations from different batches while ensuring their separation from other cell types. Visual inspection suggests that Seurat and Harmony achieve groupings mainly driven by the cell types, whereas the other methods tend to aggregate the different batches. It is notable that BBKNN brings cell populations closer but is unable to superimpose the batches.

**Figure 2.**
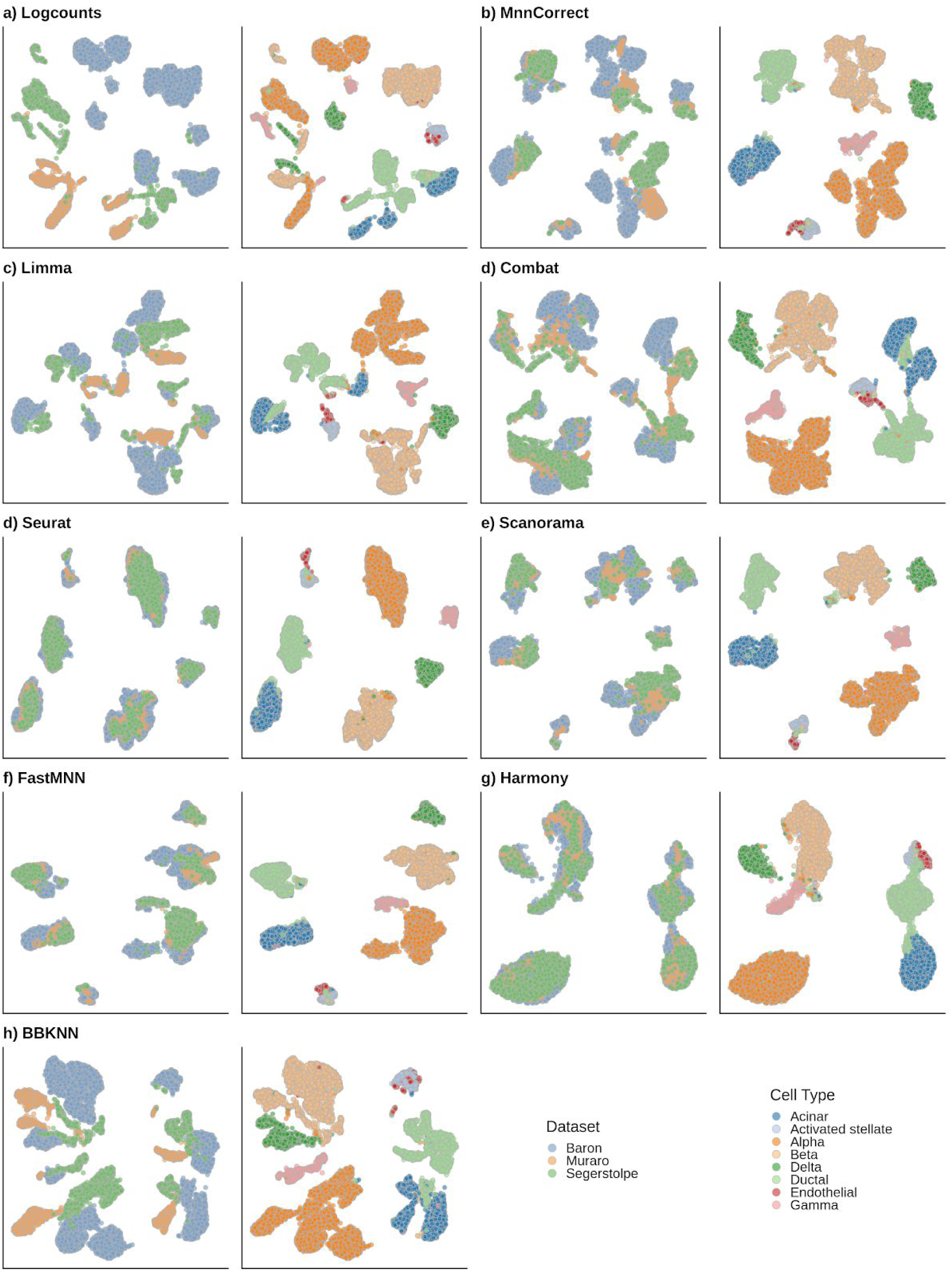
UMAP visualization of the different batch effect correction methods for the human pancreas dataset. (a-h) Each pair of panels shows the cells labeled either by dataset of origin (left) or cell type (right).

To evaluate how well the batch correction methods mix cells from different batches while keeping cell types separate, we computed the normalized Shannon entropy (16,29) based on the batch and cell type annotations provided by the original authors (Methods). The desired outcome is a high batch entropy, indicating a homogeneous mixture of the batches, and a low cell type entropy, suggesting that cell populations remain distinct. While all the methods were able to keep the distinct cell populations separate, we observed greater differences for the batch entropy (Fig 3). Based on this metric we consider Seurat and Harmony as the best methods. As intermediate performers Scanorama and fastMNN show a wider distribution of batch entropy values. Finally, mnnCorrect, Limma and ComBat can be considered the poorer performers in aligning the different batches.

**Figure 3.**
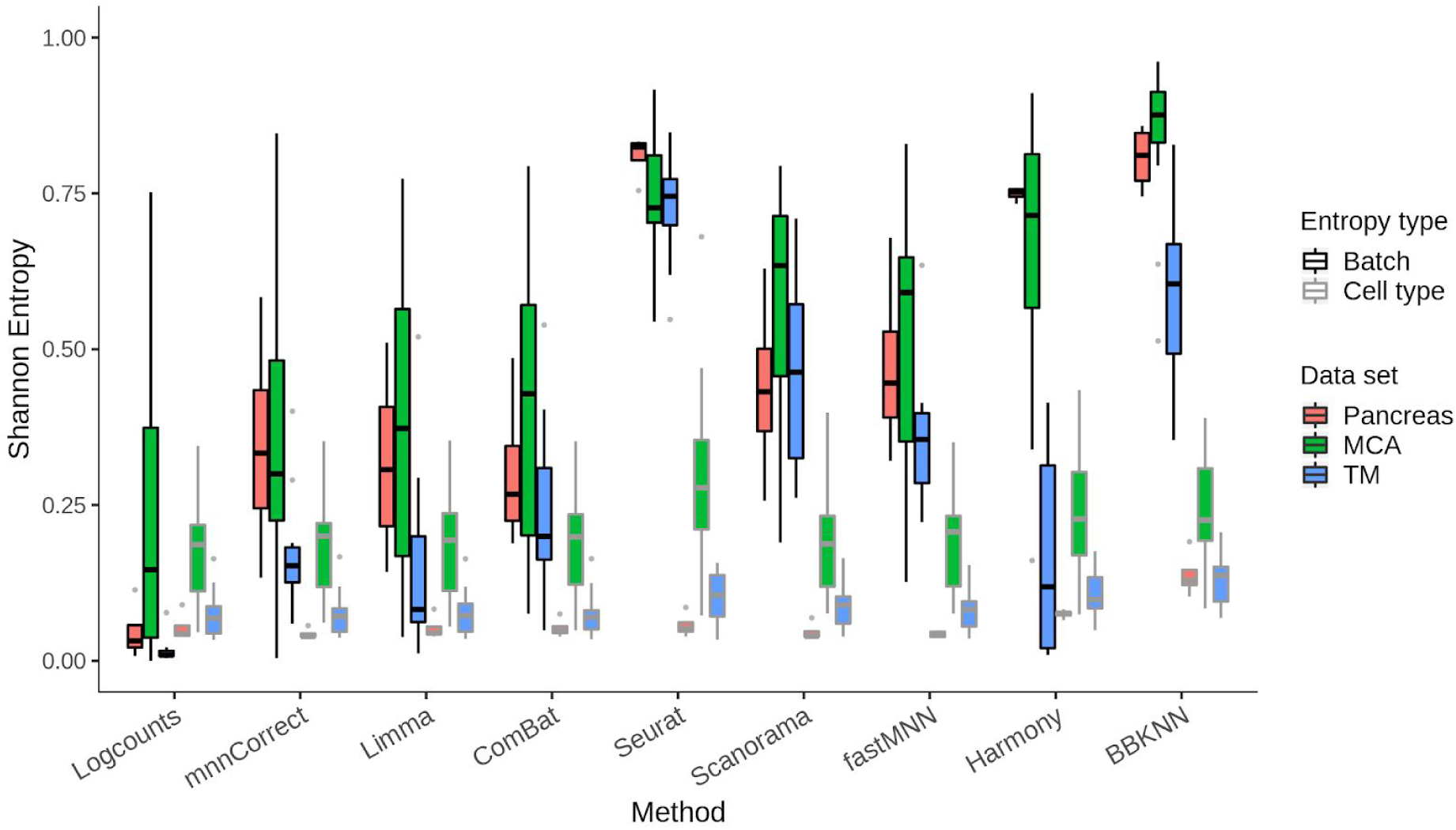
Batch and cell type entropies for eight methods and three datasets. The boxplots show the Shannon entropy over batch and cell type of the different batch effect correction methods for pancreas data (red), Mouse Cell Atlas (green), and Tabula Muris (blue). The black line represents the mean across the cells, the box the upper and lower quartiles, the whiskers 95th percentiles and the dots show outliers.

We carried out similar investigations for the Mouse Cell Atlas (MCA) (31) and Tabula Muris (32) datasets. In the MCA the batches correspond to the eight different animals (31), and as the mice all come from the same genetic background and were raised in the same environment we expect the batch effects to be smaller than for the pancreas data. The batch entropy for the uncorrected data is indeed higher than for the pancreas data (Fig 3), and most methods are able to mix the batches of the MCA better, as confirmed by visual inspection. The cell type entropies are higher than for the pancreas data, and we hypothesize that this is a consequence of the fine-grained annotation which makes it difficult to separate cell types. For example, the bone marrow contains six different types of neutrophils and the testes five types of spermatocytes. Overall across MCA data, Seurat and Harmony show the best batch mixing, although at the cost of slightly increasing cell type mixing compared to the uncorrected counts and the other methods. Scanorama can also be considered a good performer followed by fastMNN.

Next, we investigated another mouse cell atlas, Tabula Muris (32), and our analysis shows a greater sample effect as evidenced by a very low batch entropy for the uncorrected data (Fig 3). Since the batches correspond to two different experimental platforms (32), it is not surprising that there are larger differences than for the MCA. Furthermore, all methods perform better with regards to the cell type entropy, potentially due to a more coherent annotation. For all three datasets, we note that for most methods there is greater variation in batch entropy than cell type entropy. Closer inspection reveals that the batch entropies vary substantially across tissues (Table S1). Interestingly, all methods, except for Seurat and BBKNN, are unable to achieve high batch entropy for datasets with a small number of cell types. Closer inspection reveals that all methods except Seurat and BBKNN show a significant correlation between cell type entropy and number of cell types, suggesting poorer performance with more fine-grained annotation (Fig S1). Taken together, Seurat consistently succeeds in mixing the batches, again at the cost of a slightly distinct cell population mixing. Scanorama performs well although with higher variation across datasets. Surprisingly, Harmony is unable to properly align the Tabula Muris batches.

### Batch correction becomes harder as the number of cells and the number of batches increase

To determine how the number of cells in each sample influences batch correction performance and running times we considered the Tabula Muris dataset, and downsampled it to 1%, 5%, 10%, 20% and 50% of the original 60,828 cells (Methods). Across all subsets, the input objects contain 64% of 10X cells and 36% of FACS-sorted Smart-Seq2 cells. Note that this batch correction task is more challenging than the one in figure 3 as we now merge cells from different tissues.

The number of cells has a strong impact on performance and it becomes more difficult to align the two batches with increasing cell numbers. All methods except Scanorama, Harmony and Seurat reduce the batch entropy by >50% as the number of cells increases from 608 to 60,828 (Fig. 4a). Unfortunately, Scanorama mixes the cell types as well as batches, and surprisingly none of the entropies change as the number of cells increases. Harmony is the only method that, after an initial drop, increases the batch entropy with the number of cells. For all methods except Scanorama, the cell type entropy is also reduced, suggesting that it becomes easier to group cells from the same origin for larger datasets. With the exception of Scanorama, the majority of the methods do not significantly increase the cell type entropy above the value of the uncorrected counts, even decreasing it for the smaller subsets.

**Figure 4:**
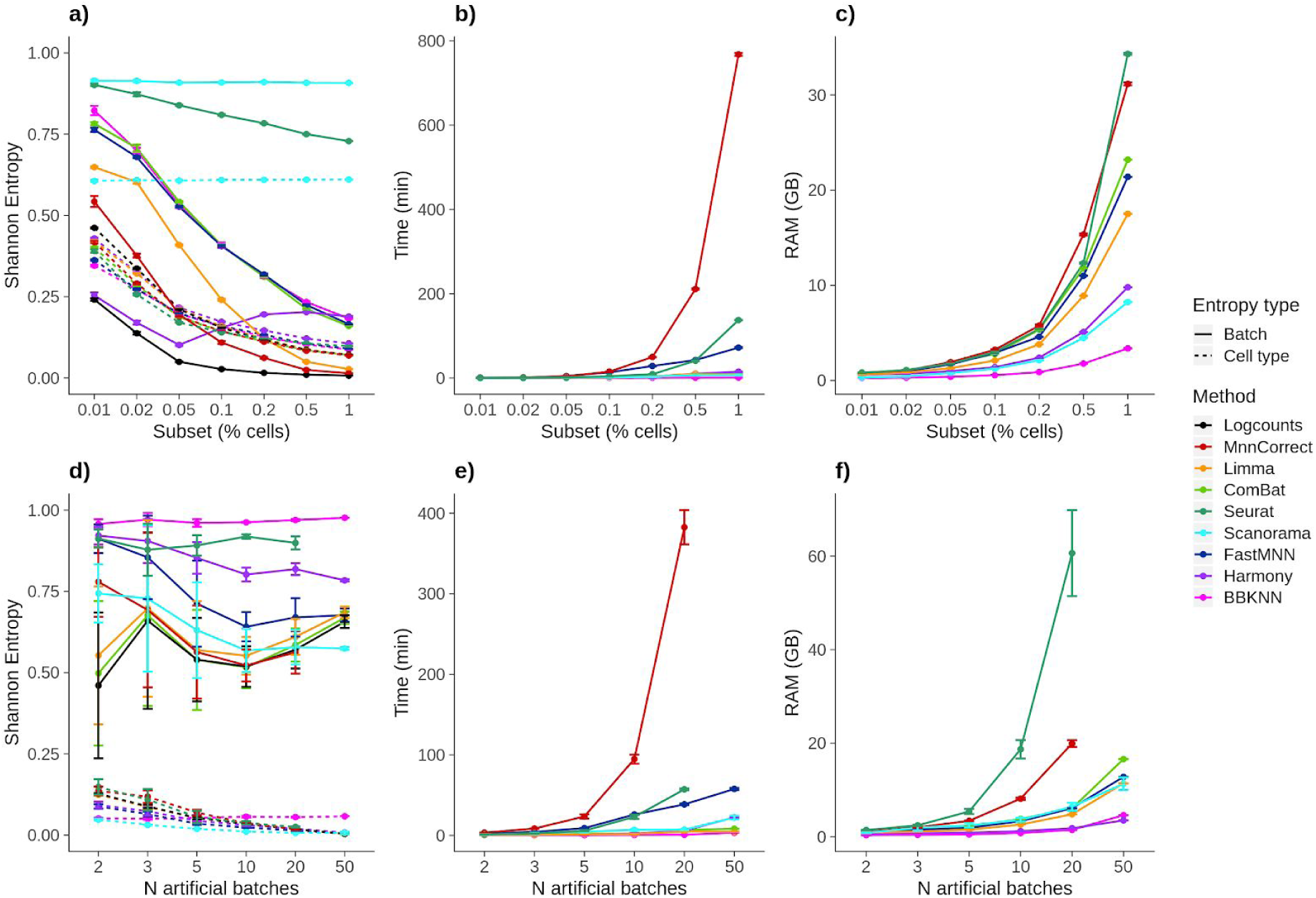
Performance of methods as a function of the number of cells and batches. a,d) Entropy, b,e) running time and c,f) RAM usage for the Tabula Muris subsets of different sizes and different numbers of simulated batches.

The main goal of the investigation involving different numbers of cells is to learn how the computational resource requirements change as this is an important factor when choosing a method. Considering the time required to perform the integration, we found substantial differences as ComBat, Limma, Harmony and BBKNN have more or less constant run times as the number of cells grow. By contrast, mnnCorrect and fastMNN grow exponentially, with the former being the slowest method in our study. Seurat initially has a stable runtime before it starts to grow exponentially (Fig 4b). For all methods we found that memory usage increases exponentially with the number of cells. The differences are smaller than for the run-time, with Seurat, mnnCorrect, ComBat and fastMNN consuming the most resources, while Harmony, Scanorama and BBKNN have the lowest requirements (Fig 4c). The memory requirements and runtimes observed in the scaling experiments are similar to what we found for the previous section (Fig S2).

As sequencing costs decrease, the number of different samples that can be processed will increase. Thus, we also evaluated how well each method handles an increasing number of batches. For this study we considered subsets of the Tabula Muris 10X dataset with 4,168 genes and 18,347 cells. As the batches created by subsampling this dataset are entirely artificial, we added small batch-specific random counts to each gene to ensure that there are differences that require correction (Methods). In our simulations, cell types are well separated whereas the batches are more overlapping.

We fixed the batch size to 1,001 cells and we created datasets including 2, 3, 5, 10, 20, and 50 and batches, introducing small artificial batch effects. Cell type entropies are maintained low with the number of batches for all methods, highlighting the capacity of our batch simulating procedure to not mix distinct cell populations as batches are included. Regarding batch entropy (Fig. 4d), BBKNN, Seurat and Harmony show the most stable performance as the number of batches increases. Although all methods have an exponential increase in both memory use and runtime, mnnCorrect stands out again as the slowest method. As before, we find that Seurat consumes the most memory, and along with mnnCorrect it fails to integrate 50 batches.

### Impact of batch correction on unsupervised clustering and identification of marker genes

A key advantage of the entropy measures is that they can easily be calculated for any dataset containing discrete cell state clusters and that they are easy to interpret. However, they only evaluate the mixing of the cells as represented by the nearest neighbor graph, and they do not directly assess how the batch correction will impact downstream analyses based on the corrected data. To understand how specific aspects of the analysis are affected, tailored benchmarks are required. BatchBench allows users to add customized modules to evaluate the aspect they find most relevant. Here, we consider two common types of analyses, unsupervised clustering and identification of marker genes.

To evaluate the effect on unsupervised clustering, we apply three popular methods, Leiden (33), Louvain (34) and SC3 (35), to the corrected data, and we then compare the merged cluster labels to the ones that were assigned prior to merging. To assess the proximity between clusterings we used a distance metric, variation of information, and a similarity metric, Adjusted Rand Index (ARI). The two measurements are by definition inversely correlated, and because they are consistent (Spearman’s rho = -.87) we will mainly refer to the ARI results.

Our analysis of the MCA suggested small differences in cell type entropy, but large differences in how well the batches were mixed (Fig 3). By contrast, when running unsupervised clustering the batch correction methods achieve similar ARI values, with only small differences between the Louvain, Leiden and SC3 algorithms (Fig 5a, S3-6). Closer inspection instead reveals large differences between tissues, something that is not evident from the entropy measures (Table S1). For the Tabula Muris we observe a similar pattern with large differences in ARI between tissues and relatively small differences across methods. The main difference compared to the MCA is that the clusters reported by SC3 have a higher ARI than the ones reported by the Louvain and Leiden methods for 7 of 11 tissues. Closer inspection reveals that the Leiden and Louvain methods perform poorly for datasets with a small number of clusters (Fig. S3,S4). Surprisingly, for heart and mammary glands, the best clustering results are achieved with SC3 applied to the uncorrected data. For the pancreas datasets we find that SC3 tends to have a higher ARI, and unlike the two mouse atlases there is good agreement with the entropy analysis as Seurat and Harmony performed the best.

**Figure 5:**
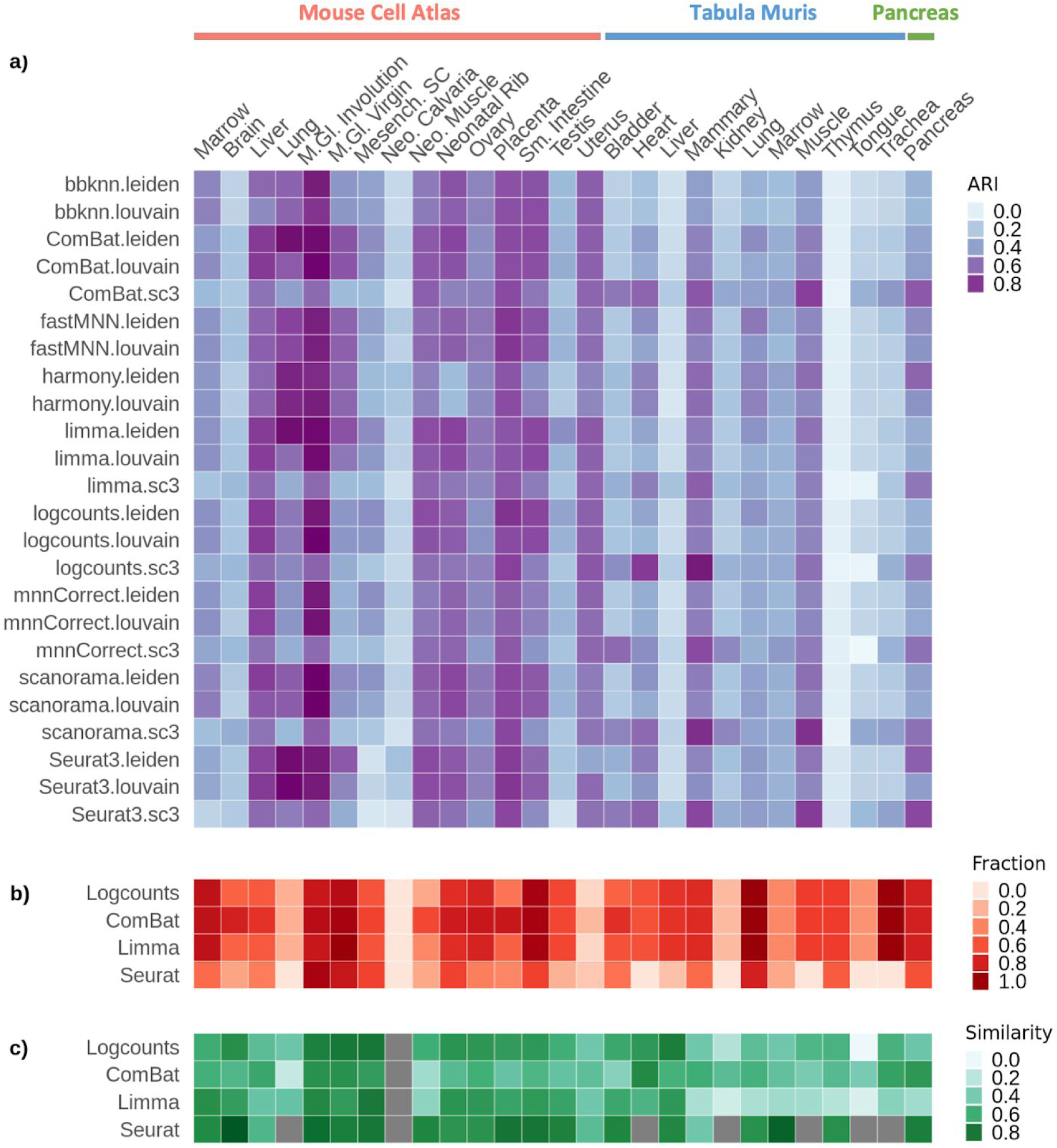
Evaluation of the impact of batch correction on unsupervised clustering and marker gene identification. a) Clustering similarity of batch corrected output to cell labels as evaluated by the Adjusted Rand Index. b) Fraction of total cell types over which marker genes are detected. c) Similarity of marker genes as evaluated by the generalized Jaccard Index.

The main objective of batch correction methods is to ensure that cells with similar expression profiles end up near each other. The most widely used metrics, e.g. mixing entropies or inverse Simpson index (16,19, 29), are designed to evaluate this aspect. However, if a researcher is interested in analyzing the expression values for other purposes then it is important to make sure that the corrected values are close to the original ones. To investigate how much expression matrices are distorted by the different methods, we compared the marker genes identified before and after batch correction for the five methods that modify the expression matrix (Table 1). We identified marker genes for each batch individually as well as for the merged datasets from each method that outputs a modified expression matrix. Unlike the entropy and clustering analyses, we observed stark differences between batch correction methods. Remarkably, after merging using Scanorama or mnnCorrect, not a single marker gene is identified. Only ComBat and Limma are able to identify marker genes for most cell types, while Seurat only reports markers for a minority of cell types in most tissues (Fig 5b). Comparing the similarity between the marker genes identified in the individual batches and the merged dataset using a generalized Jaccard index (36), we find that Seurat provides the highest degree of consistency (Fig 5c). However, it is important to keep in mind that Seurat’s good performance is biased by the fact that it reports marker genes for fewer cell types than the other methods. A similar problem stems from the fact that sometimes the individual batches do not share any or only few marker genes prior to merging, e.g. the neonatal calvaria from the MCA, which explains the grey boxes in figure 5c.

## Discussion

We have developed BatchBench, a customizable pipeline for comparing scRNA-seq batch correction methods. We have assessed the performance of eight popular batch correction methods based on entropy measurements across three datasets, suffering from donor and platform effects. Our results highlight Seurat as the top performer as it correctly merges batches while maintaining the separation of distinct cell populations. Harmony also shows very good results in pancreas and MCA but surprisingly fails in correcting the Tabula Muris batch effects. Scanorama and fastMNN can be considered consistent good performers. Regarding BBKNN, we note that the entropies are not suitable for evaluating its performance as the method operates by identifying nearest neighbours in each of the provided batches (26) and adjusting neighbors to maximize the batch entropy. Hence, a different metric should be established to evaluate the performance of BBKNN. We also evaluated how the methods perform as the number of cells and the number of batches are varied. Here, we highlight Harmony as a method that provides good performance while being economical in its use of computational resources. However, our analyses suggest that all methods, with the possible exceptions of BBKNN and Harmony, will struggle to integrate hundreds of batches even if each batch is relatively small. Thus, improving scalability is a central requirement for future methods.

A key insight from our study is that the entropy measures do not fully reflect how the choice of batch correction method will impact downstream computational analyses. We applied three different unsupervised clustering methods to the merged datasets, and the results are not as clear as for the entropy analyses. No single method emerges as the best performer, and in some cases the best results were obtained using the uncorrected data. This result highlights the importance of using benchmarks that are more closely linked to the analysis that will be carried out for the merged dataset.

Our attempt to identify marker genes from the corrected dataset demonstrates the difficulty of using the merged expression matrix for downstream analyses. As none of the methods considered in our study performed adequately in this benchmark, we highlight this as an area where improvements are required. Since marker genes are not preserved, we stress the importance for users to monitor how expression levels change. Any analysis based on the expression levels, e.g. identification of marker genes or differentially expressed genes, will need to be verified to ensure that the result was not distorted due to the alterations introduced by the batch correction method. An important limitation of our marker gene analysis is that it only quantifies consistency as there is not yet an established ground truth for what marker genes are represented for the cell types in our study. We tried to use marker gene lists from the literature as represented by the CellMarker database (37), but we found that all pancrease datasets provided poor overlap, even before clustering (Fig S7).

Benchmark studies are important as they help guide researchers in their choice of methods. They are also helpful for developers as they can highlight limitations of existing methods and provide guidance as to where improvements are needed. One shortcoming of traditional benchmarks, however, is that they are static in nature and that they only consider the datasets that the authors of the benchmark study had chosen to include. A related issue is that the metrics used to evaluate methods may not be relevant to all datasets and research questions. Along with a similar study by Leucken et al (40), BatchBench will serve as a useful platform to the community as it enables benchmarks to be tailored to specific needs.

## Methods

### Datasets

#### Pancreas dataset

We consider three published pancreas datasets: Baron (<u>GSE84133</u>) (39), Muraro (<u>GSE85241</u>) (27), and Segerstolpe (<u>E-MTAB-5061</u>) (28) generated using inDrop, CEL-Seq2 and Smart-Seq2 technologies, respectively. Initially, quality control was performed on each of the datasets to remove cells with <200 counts and genes that were present in <3 cells along with spike-ins and anti-sense transcripts. Furthermore, we only retained cells that had been assigned a biologically meaningful cell type (e.g. removing cells from the “unclassified” category).

For figure 3 we wanted to represent the pancreas results as a boxplot similar to the other datasets. To ensure that we got a distribution we considered three additional versions of the data. One of these versions contained all of the genes expressed across the three batches rather than just the highly variable ones. The second contained 1,000 cells selected randomly from each batch using the highly variable genes. The third version contained only six cell types (acinar, alpha, beta, gamma, delta and ductal) from each batch downsampled to 50% of the original number of cells and information from the highly variable genes.

#### Mouse Cell Atlas datasets

Individual MCA datasets were downloaded from https://figshare.com/s/865e694ad06d5857db4b and merged by tissue, generating 37 organ datasets. From these, 18 datasets containing more than 1 batch and with a reasonable proportion of cells across batches were selected. Through further preprocessing we removed cells expressing <250 genes, genes expressed in <50 cells, cell types representing <1% of total cell population in a tissue, and batches containing <5% of the total number of cells in a tissue (Table S2).

#### Tabula Muris datasets

The data was downloaded from https://www.google.com/url?q=https://figshare.com/projects/Tabula_Muris_Transcriptomic_characterization_of_20_organs_and_tissues_from_Mus_musculus_at_single_cell_resolution/27733&sa=D&ust=1589187433512000&usg=AFQjCNFC_0CGNwum-u2nka-OvFAmxoECtA. For all analyses except figure 4, individual datasets representing the same tissue across the two platforms were merged into 11 organ datasets (Table S1). We set workflow quality control parameters to remove cells expressing <1000 genes, genes expressed in <50 cells. Again, cell types representing <1% of total cell population in a tissue, and batches containing <5% of the total number of cells in a tissue were excluded from further analyses. For the scaling analysis in Figure 4, the previous tissues were merged into an atlas Tabula Muris dataset which was filtered to retain cells with >200 genes expressed, genes expressed in >3 cells. Cells assigned to NA or unknown cell types were excluded. Cell types representing <1% of total cell population in a tissue, and batches containing <5% of the total number of cells in a tissue were excluded from further analyses. This resulted in an object of 4,168 genes and 60,828 cells (40,058 from *10X* and 20,770 from *Smart-Seq2*).

### Batch and cell type entropy

The output of each tool is transformed into a K Nearest Neighbour graph with each node *i* representing a cell (`BuildKNNGraph`, scran package). Each cell is connected to its *k*=30 nearest neighbors as defined by the similarity of expression profiles calculated using the Euclidean distance. Using the graph we calculate for each cell *i* the probability that a neighbor has cell type *c, P*_*ic*_, as well as the probability that a neighbor comes from batch *b, P*_*ib*_. From these joint probabilities we can calculate cell type and batch entropies. We report the average value across all cells divided by the theoretical maximum to ensure a value in the interval [0, 1]. For the datasets considered in this study, the results are robust with respect to the choice of *k* (Fig S8).

### UMAP

Uniform Manifold Approximation and Projection (UMAP) is computed through the `scanpy.api.tl.umap` function, which uses the implementation of umap-learn (38). For the batch removal methods implemented in R, the rds objects are first converted into h5ad objects using the `sce2anndata` from the sceasy package (https://github.com/cellgeni/sceasy/).

### Downsampling

The filtered Tabula Muris dataset was sampled using uniform selection and no replacement to 1, 2, 5, 10, 20, and 50 percent of its cells. Resulting in objects of: 4168 genes and 608, 1217, 3041, 6083, 12166, and 30414 cells. The initial proportion of the batches (0.64, 0.36) was maintained through the different subsets.

### Artificial batches

We work with a reduced version of the Tabula Muris atlas object. We first removed all the Smart-seq2 cells and then retained only the 10 largest cell types. From this 1,001 cells are randomly sampled to serve as input to the artificial batch generation. All 4,168 initial genes are considered. We base our simulation of batch effects on a normal distribution. For each batch to be simulated, we define: i) a fraction *f* of cells sampled with uniform probability from the sequence [0.05, 0.1, 0.15, 1.0]; ii) a value *d* representing the dispersion of the effect to be simulated sampled with uniform probability from the sequence [0.5, 1.0, 1.5, … *n*], where *n* is the number of batches to simulate. For each of the 10 cell types in the input data we add count values by drawing values from a normal distribution with a standard deviation *d*. The artificial batch effect is only applied to those genes expressed in >*f* of the cells. If a gene is assigned a negative value, then it is replaced by 0. The result is a simulated data set of 1,001 cells and 4168 genes which is appended to the input data set. We followed this approach to simulate data sets with 2, 3, 5, 10, 20 and 50 equally sized batches.

### Clustering analysis

The merged samples were clustered using SC3 (35) from the homonim Bioconductor package, as well as the Louvain and Leiden algorithms implemented in Seurat (23). SC3 requires a count matrix as input, whereas Seurat can operate on a low dimensional representation. For SC3 we set *k* to the number of cell populations of each dataset. If the dataset had >5,000 cells we enable `sc3_run_svm` to speed up the processing. For the other methods we used the Seurat function `FindClusters`, specifying Louvain original algorithm and Leiden algorithm, with other parameters set to their default values.

### Marker Gene Analysis

To obtain marker genes we use the `FindMarkers` function from the Seurat package which restricts the comparison to methods that output a normalised count matrix. For a gene to be considered as a marker, we require that the absolute value of the log fold-change >2, and that the gene is expressed in at least half of the cells in each population. We use the default Wilcoxon Rank Sum test to find genes that are significantly different (adjusted p-value<0.05) between the merged dataset, and in each of the individual batches.

To compare the overlap of the sets of marker genes identified across batches and the merged data we used the multiple site generalized Jaccard index (36). We restricted the comparison to the cell populations that are common to all individual batches. We also investigate the proportion of cell populations of the dataset for which marker genes can be found.

### BatchBench pipeline

As an input, BatchBench (https://github.com/cellgeni/batchbench) requires equivalent `SingleCellExperiment` (for the R based methods) and `AnnData` objects (for the python based methods). These objects must contain: log-normalized counts, and the batch and cell type annotation of their cells as `Batch` and `cell_type1` respectively, in the object metadata. The workflow performs an initial QC step where cells, genes, batches or cell types can be filtered according to user-defined parameters. Cells not assigned to any batch or cell type are excluded in this step also. Each dataset is then sent in parallel as input to each of the batch effect correction tools, after which rds and h5ad objects containing the output are saved and made available for the user. Each of the batch corrected outputs serves as input for a series of downstream analyses: (i) UMAP coordinates are computed and saved as a csv file for visualization of the different batch corrections, (ii) Entropy computation and saved as csv file, (iii) Clustering analysis, (iv) Marker gene analysis and any module optionally added by the user.

## Author Contributions

MH and VYK conceived of the study and supervised the work. RCG and SvD developed the Nextflow pipeline. RCG carried out the benchmarking of the three datasets and eight methods. RCG and MH wrote the manuscript with inputs from VYK and SvD.

## Acknowledgements

We would like to thank members of the Cellular Genetics Informatics team and the Hemberg lab for constructive feedback and comments. The SvD, VYK and MH were funded by a core grant from the Wellcome Trust. RCG was funded by the Polytechnic University of Valencia under an Erasmus+ studentship, and by the Wellcome Trust.

## Supplementary Materials

**Figure S1.**
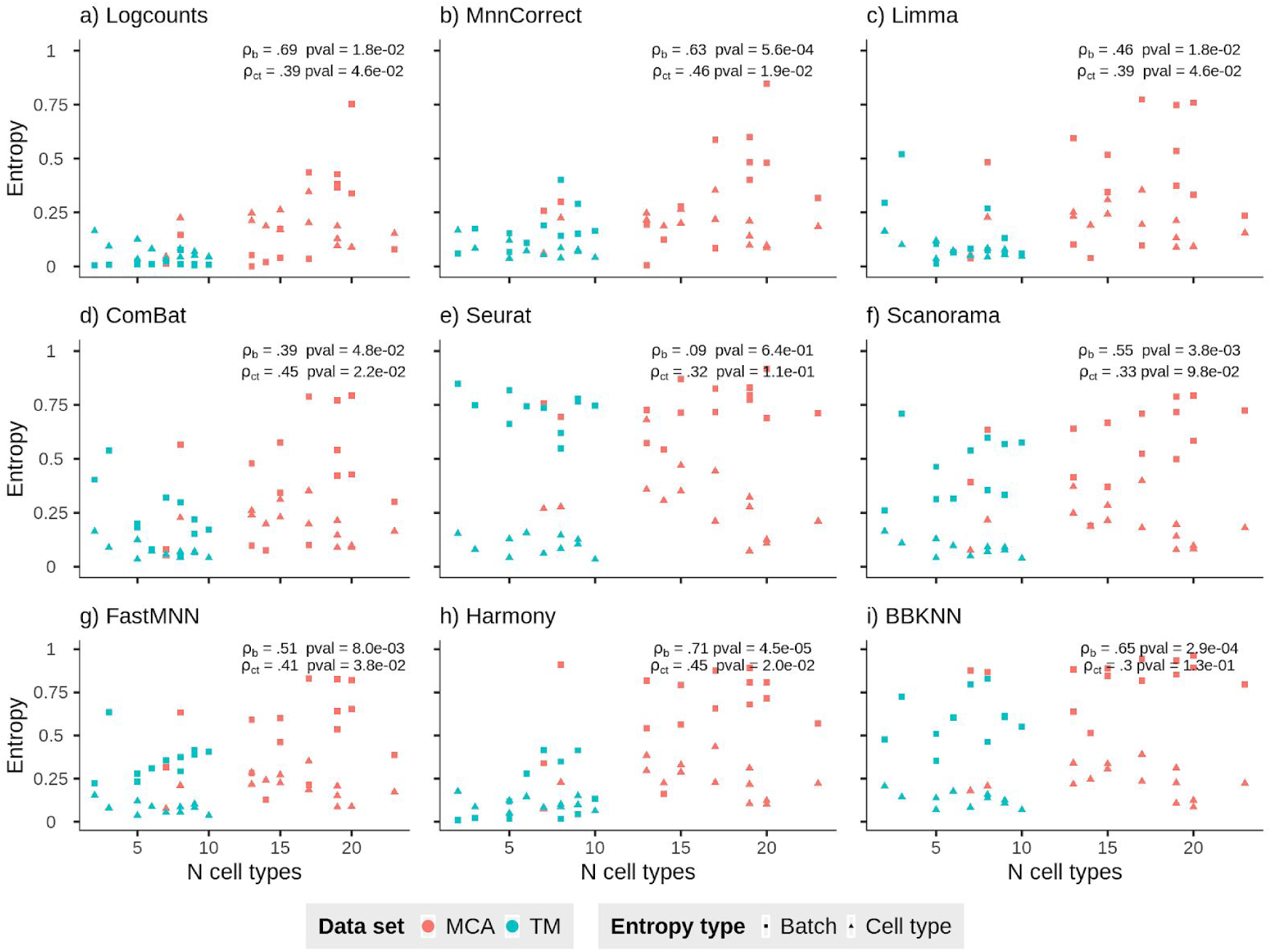
Batch and cell type entropies for eight methods and three datasets as a function of the number of cell types. The inset text for each panel shows the Spearman’s rank correlation coefficient between the number of cell types and batch (ρ_b_), or cell type (ρ_ct_) entropy values.

**Figure S2.**
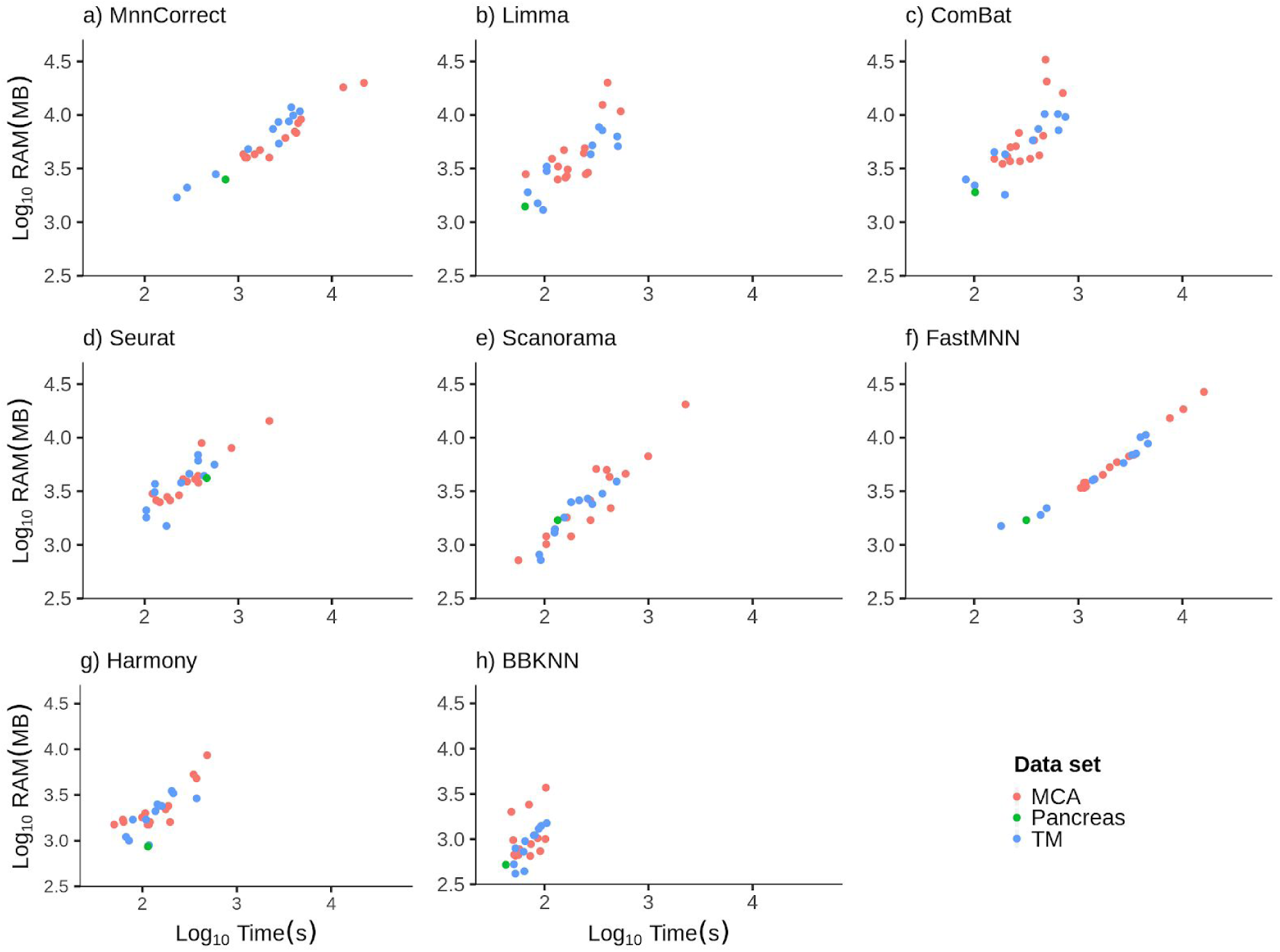
Memory requirements and runtimes for all datasets per method.

**Figure S3.**
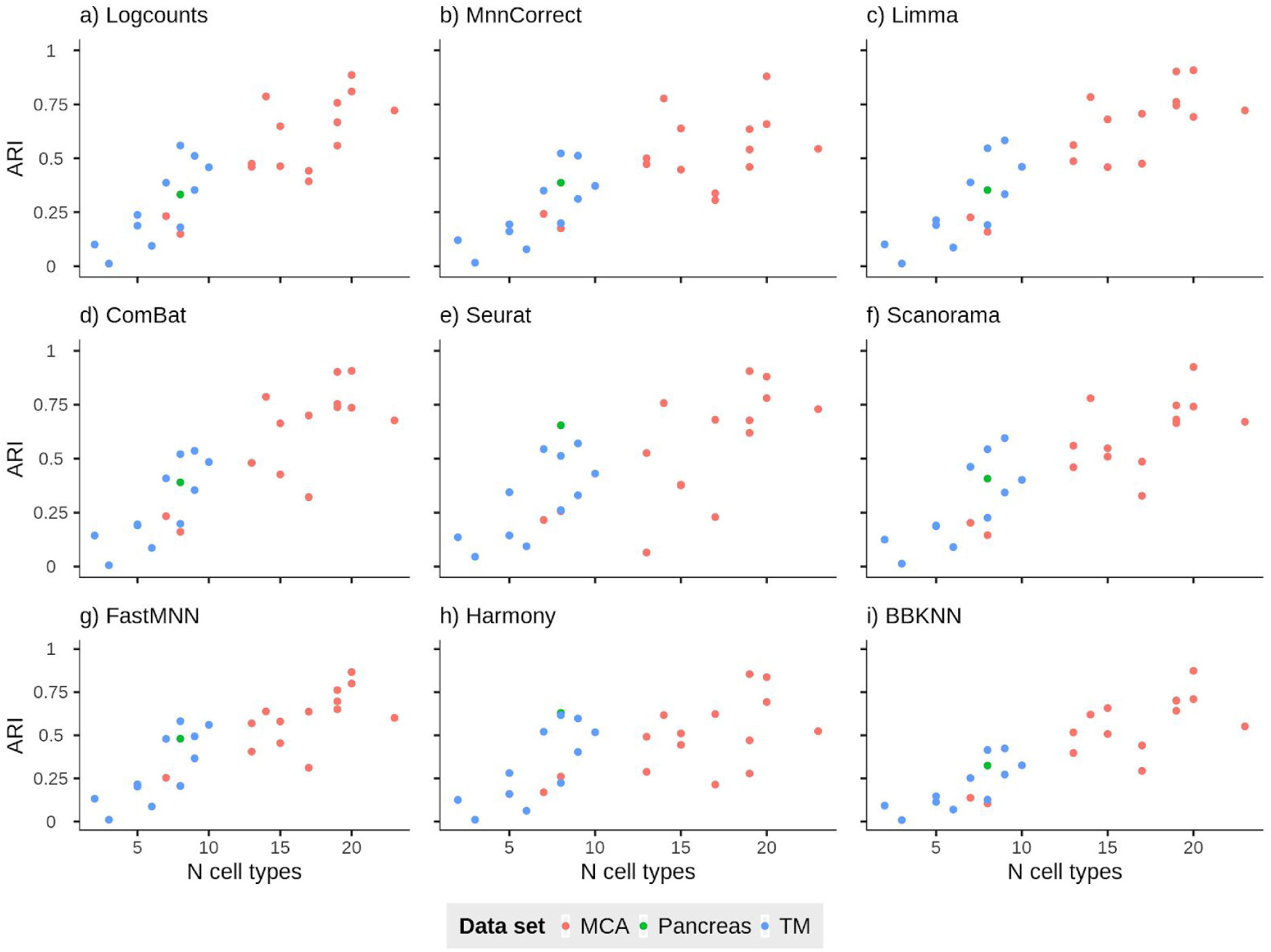
Adjusted Rand Index for the Leiden clustering algorithm as a function of the number of cell types.

**Figure S4.**
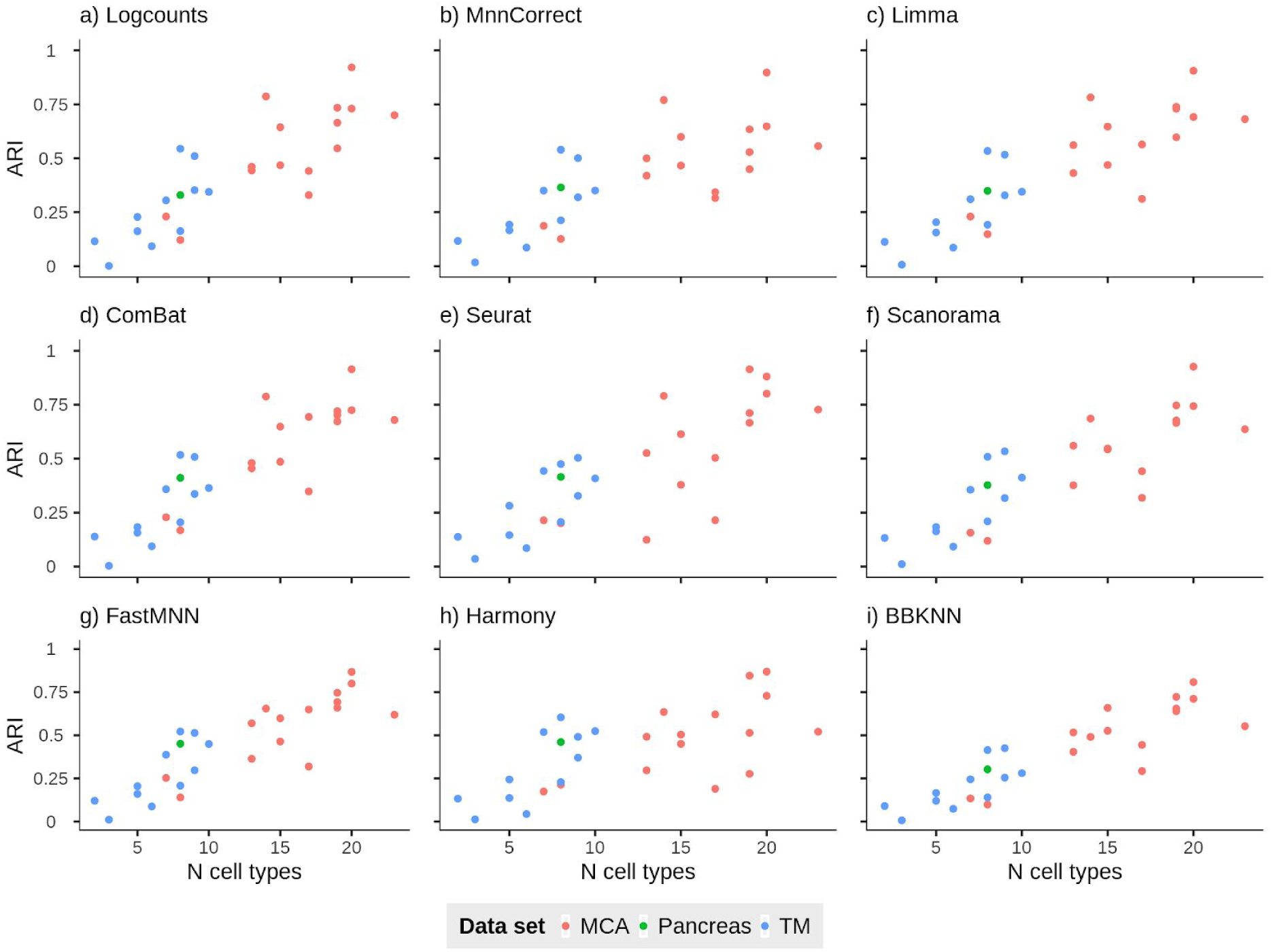
Adjusted Rand Index for the Louvain clustering algorithm as a function of the number of cell types.

**Figure S5.**
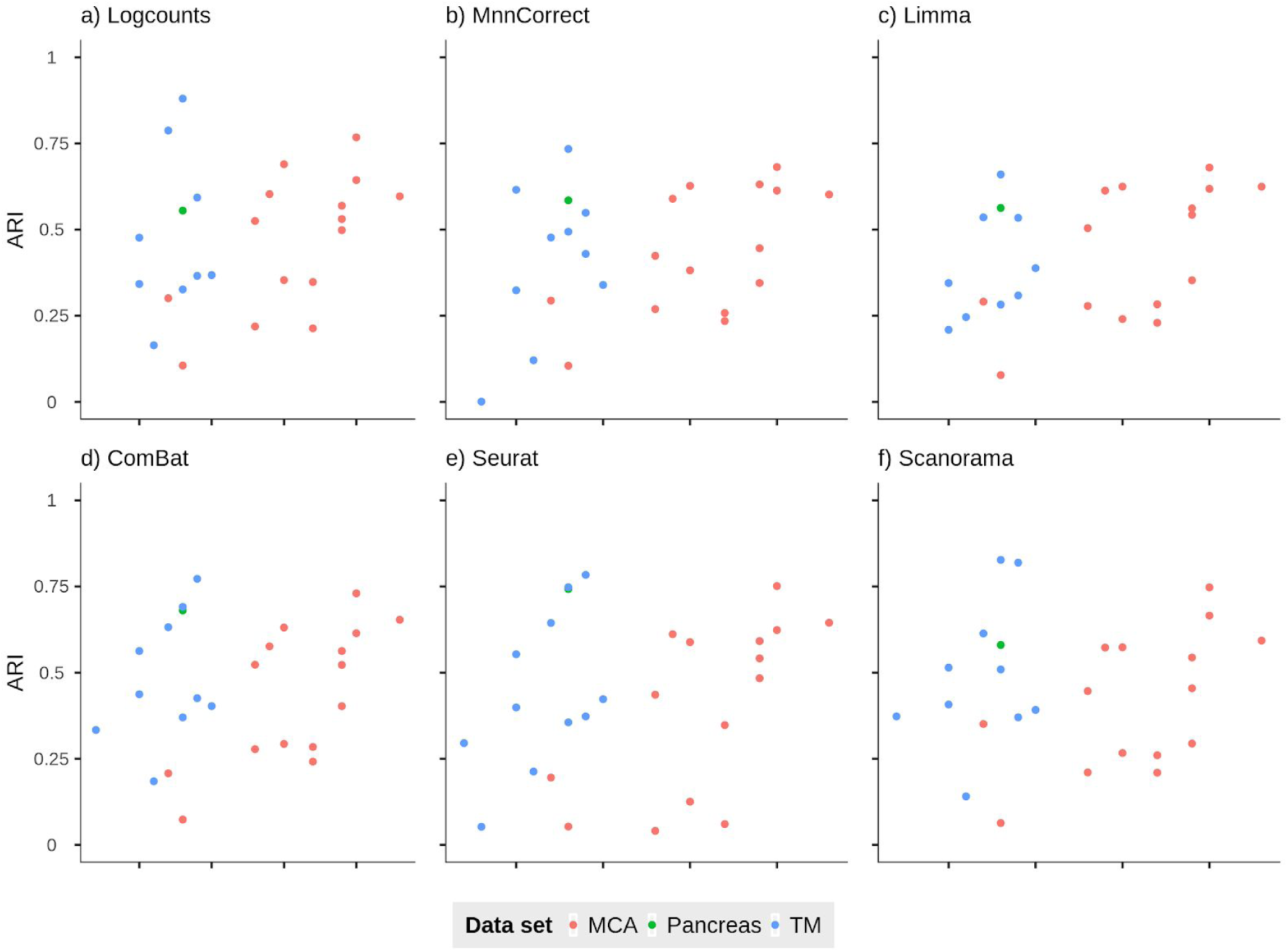
Adjusted Rand Index for the SC3 clustering algorithm as a function of the number of cell types.

**Fig S6.**
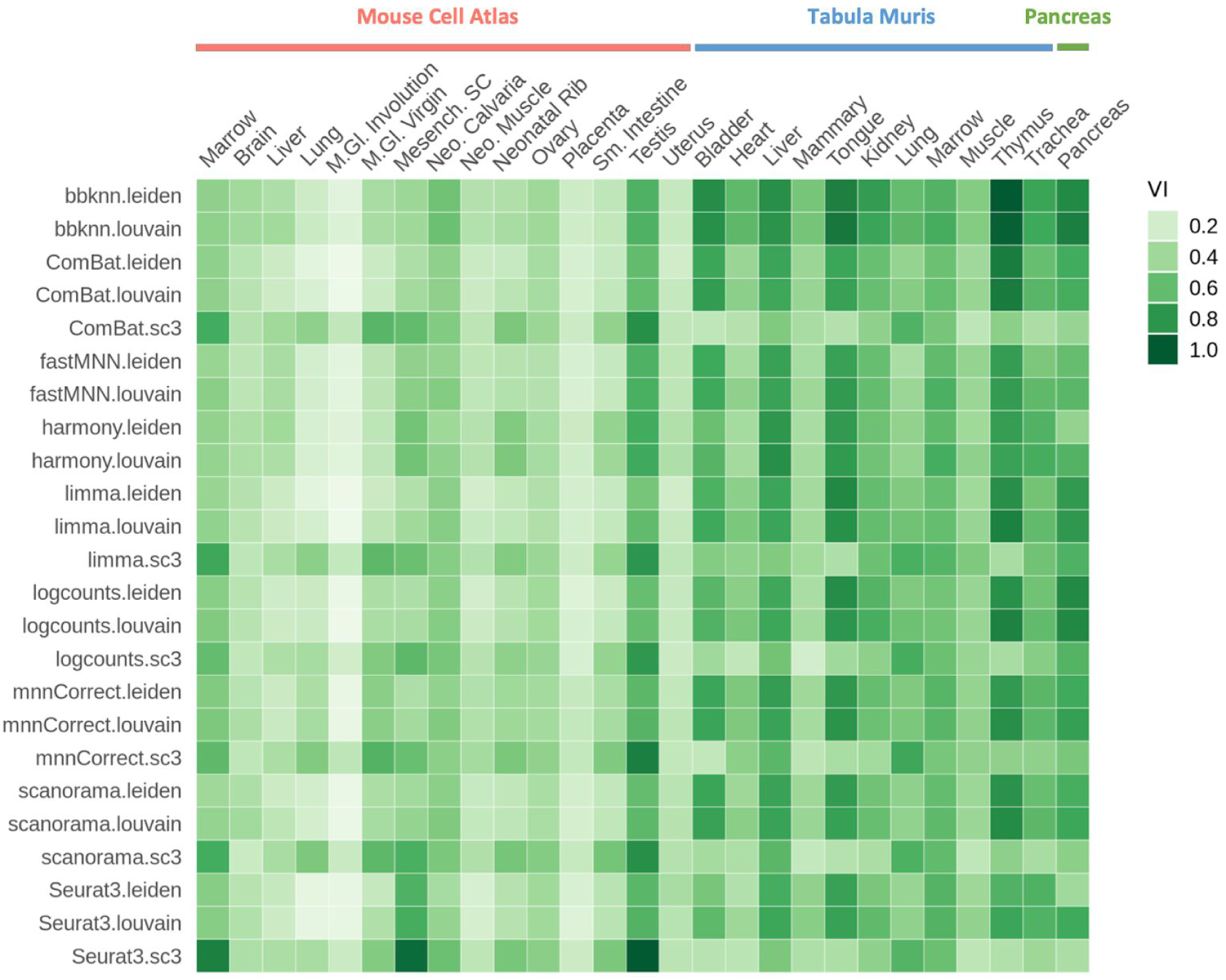
Clustering similarity of batch corrected output to cell labels as evaluated by Variation of Information distance.

**Figure S7.**
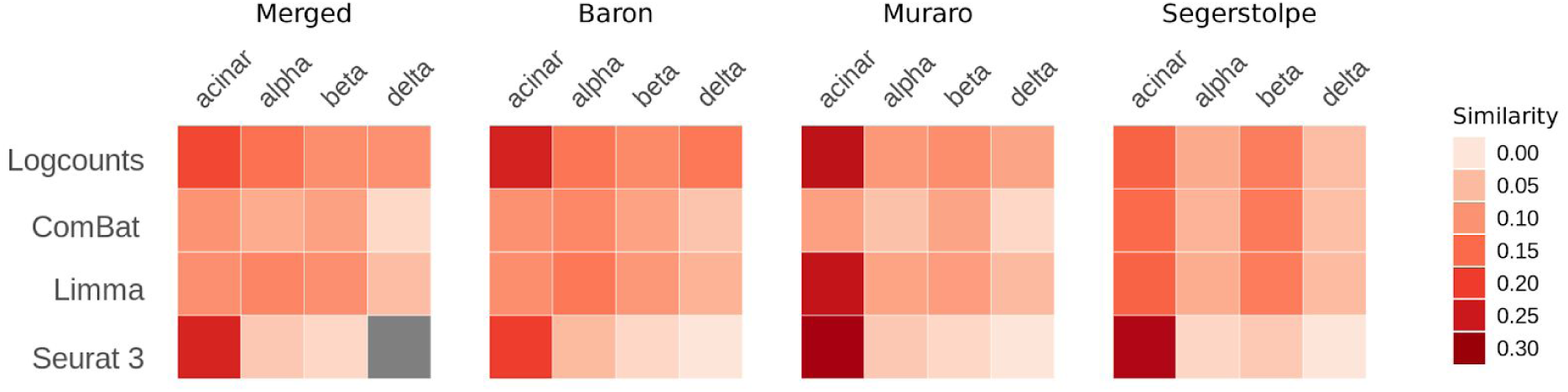
Jaccard similarity index computed between Pancreas and CellMarker database markers.

**Figure S8.**
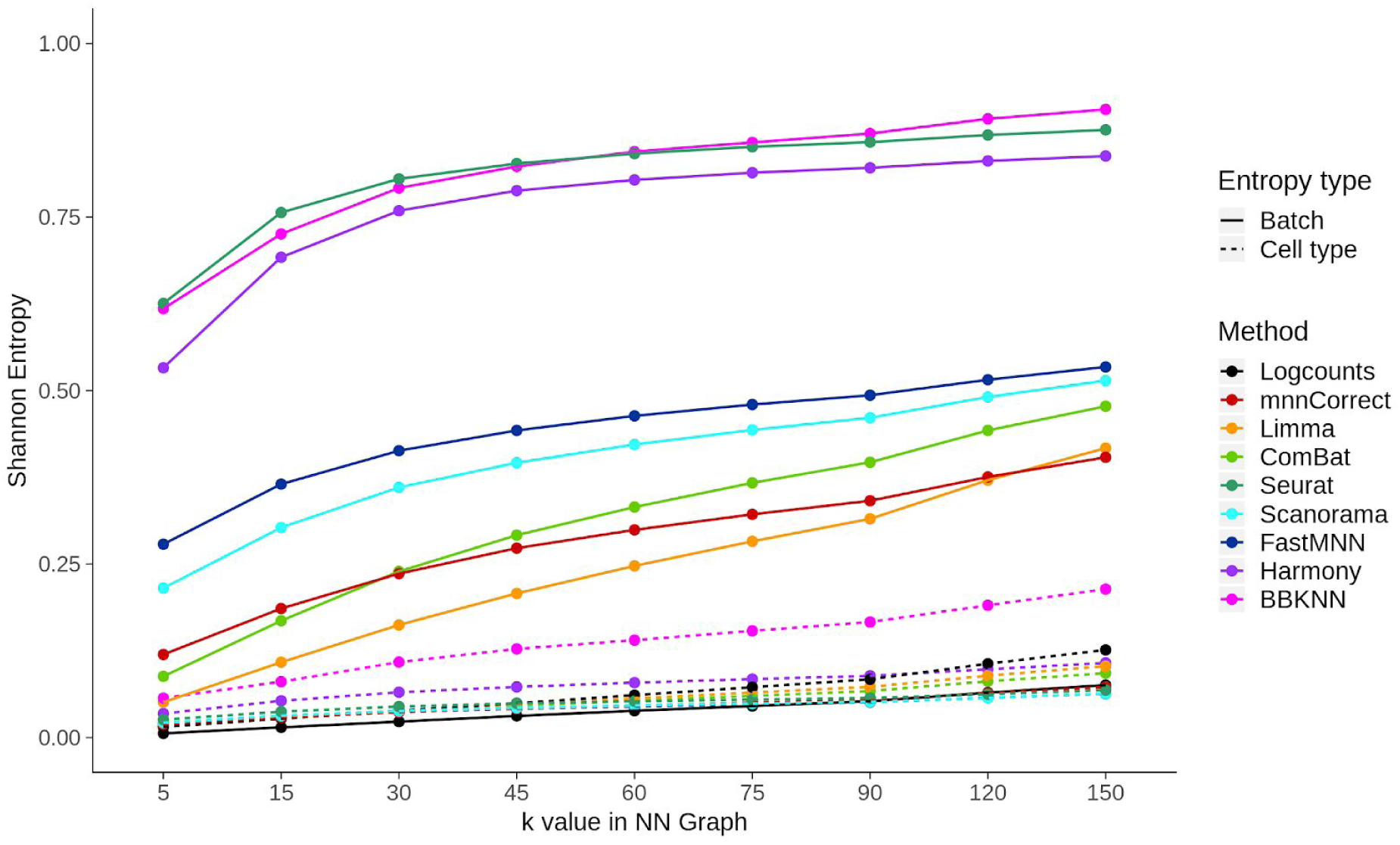
Batch and cell type entropy for the pancreas dataset using different values of k for the Pancreas dataset with cross-batch common genes.

**Table S1.** Entropy values per dataset (MCA: Mouse Cell Atlas, TM: Tabula Muris).

| N | Data set | Organ | Entropy type | Log-counts | mnn-Correct | Limma | Com-Bat | Seurat | Scano-rama | fast-MNN | Har-mony |
| --- | --- | --- | --- | --- | --- | --- | --- | --- | --- | --- | --- |
| 1 | MCA | Bone Marrow | Batch | 0.172 | 0.27 | 0.517 | 0.576 | 0.87 | 0.667 | 0.601 | 0.793 |
| 2 | MCA | Bone Marrow | Cell type | 0.262 | 0.265 | 0.308 | 0.312 | 0.47 | 0.284 | 0.272 | 0.33 |
| 3 | MCA | Brain | Batch | 0.011 | 0.257 | 0.038 | 0.08 | 0.757 | 0.393 | 0.316 | 0.339 |
| 4 | MCA | Brain | Cell type | 0.046 | 0.061 | 0.055 | 0.049 | 0.269 | 0.076 | 0.076 | 0.075 |
| 5 | MCA | Liver | Batch | 0.02 | 0.125 | 0.039 | 0.076 | 0.544 | 0.19 | 0.126 | 0.161 |
| 6 | MCA | Liver | Cell type | 0.186 | 0.187 | 0.19 | 0.199 | 0.307 | 0.188 | 0.241 | 0.224 |
| 7 | MCA | Lung | Batch | 0.427 | 0.598 | 0.748 | 0.771 | 0.797 | 0.788 | 0.827 | 0.892 |
| 8 | MCA | Lung | Cell type | 0.095 | 0.097 | 0.088 | 0.089 | 0.073 | 0.078 | 0.085 | 0.104 |
| 9 | MCA | M. Gl. Involution | Batch | 0.752 | 0.846 | 0.759 | 0.794 | 0.917 | 0.794 | 0.819 | 0.809 |
| 10 | MCA | M. Gl. Involution | Cell type | 0.089 | 0.087 | 0.09 | 0.09 | 0.109 | 0.083 | 0.087 | 0.101 |
| 11 | MCA | M. Gl. Virgin | Batch | 0.437 | 0.587 | 0.774 | 0.787 | 0.825 | 0.71 | 0.83 | 0.877 |
| 12 | MCA | M. Gl. Virgin | Cell type | 0.202 | 0.218 | 0.194 | 0.198 | 0.211 | 0.18 | 0.185 | 0.228 |
| 13 | MCA | Mesench. SC | Batch | 0 | 0.004 | 0.101 | 0.098 | 0.574 | 0.415 | 0.282 | 0.541 |
| 14 | MCA | Mesench. SC | Cell type | 0.211 | 0.215 | 0.231 | 0.24 | 0.68 | 0.371 | 0.285 | 0.384 |
| 15 | MCA | Neonatal Calvaria | Batch | 0.146 | 0.3 | 0.482 | 0.566 | 0.695 | 0.634 | 0.633 | 0.911 |
| 16 | MCA | Neonatal Calvaria | Cell type | 0.225 | 0.224 | 0.227 | 0.227 | 0.278 | 0.217 | 0.21 | 0.228 |
| 17 | MCA | Neonatal Muscle | Batch | 0.08 | 0.317 | 0.235 | 0.301 | 0.711 | 0.723 | 0.388 | 0.57 |
| 18 | MCA | Neonatal Muscle | Cell type | 0.152 | 0.184 | 0.155 | 0.165 | 0.211 | 0.18 | 0.172 | 0.221 |
| 19 | MCA | Neonatal Rib | Batch | 0.381 | 0.484 | 0.534 | 0.542 | 0.773 | 0.717 | 0.64 | 0.809 |
| 20 | MCA | Neonatal Rib | Cell type | 0.187 | 0.21 | 0.211 | 0.214 | 0.322 | 0.197 | 0.207 | 0.31 |
| 21 | MCA | Ovary | Batch | 0.052 | 0.193 | 0.595 | 0.479 | 0.727 | 0.639 | 0.591 | 0.816 |
| 22 | MCA | Ovary | Cell type | 0.247 | 0.247 | 0.25 | 0.261 | 0.358 | 0.249 | 0.217 | 0.296 |
| 23 | MCA | Placenta | Batch | 0.338 | 0.48 | 0.332 | 0.428 | 0.69 | 0.584 | 0.655 | 0.715 |
| 24 | MCA | Placenta | Cell type | 0.089 | 0.097 | 0.093 | 0.098 | 0.125 | 0.097 | 0.088 | 0.122 |
| 25 | MCA | Small Intestine | Batch | 0.366 | 0.402 | 0.373 | 0.421 | 0.83 | 0.498 | 0.534 | 0.679 |
| 26 | MCA | Small Intestine | Cell type | 0.128 | 0.14 | 0.132 | 0.147 | 0.277 | 0.141 | 0.151 | 0.217 |
| 27 | MCA | Testis | Batch | 0.035 | 0.084 | 0.097 | 0.101 | 0.716 | 0.524 | 0.215 | 0.656 |
| 28 | MCA | Testis | Cell type | 0.345 | 0.352 | 0.353 | 0.352 | 0.443 | 0.398 | 0.351 | 0.435 |
| 29 | MCA | Uterus | Batch | 0.04 | 0.277 | 0.345 | 0.343 | 0.715 | 0.371 | 0.461 | 0.563 |
| 30 | MCA | Uterus | Cell type | 0.171 | 0.2 | 0.242 | 0.23 | 0.351 | 0.214 | 0.225 | 0.288 |
| 31 | TM | Bladder | Batch | 0.022 | 0.068 | 0.012 | 0.183 | 0.817 | 0.313 | 0.278 | 0.018 |
| 32 | TM | Bladder | Cell type | 0.126 | 0.119 | 0.119 | 0.125 | 0.129 | 0.13 | 0.119 | 0.123 |
| 33 | TM | Heart | Batch | 0.021 | 0.189 | 0.082 | 0.321 | 0.737 | 0.538 | 0.355 | 0.414 |
| 34 | TM | Heart | Cell type | 0.035 | 0.053 | 0.049 | 0.058 | 0.062 | 0.05 | 0.055 | 0.083 |
| 35 | TM | Kidney | Batch | 0.077 | 0.4 | 0.269 | 0.298 | 0.548 | 0.597 | 0.373 | 0.349 |
| 36 | TM | Kidney | Cell type | 0.082 | 0.085 | 0.083 | 0.07 | 0.084 | 0.091 | 0.085 | 0.086 |
| 37 | TM | Liver | Batch | 0.011 | 0.11 | 0.065 | 0.081 | 0.744 | 0.316 | 0.309 | 0.279 |
| 38 | TM | Liver | Cell type | 0.081 | 0.071 | 0.073 | 0.073 | 0.157 | 0.097 | 0.088 | 0.145 |
| 39 | TM | Lung | Batch | 0.008 | 0.165 | 0.06 | 0.171 | 0.745 | 0.576 | 0.406 | 0.133 |
| 40 | TM | Lung | Cell type | 0.045 | 0.041 | 0.045 | 0.041 | 0.034 | 0.039 | 0.036 | 0.063 |
| 41 | TM | Mammary | Batch | 0.009 | 0.142 | 0.074 | 0.049 | 0.619 | 0.356 | 0.293 | 0.017 |
| 42 | TM | Mammary | Cell type | 0.043 | 0.038 | 0.043 | 0.043 | 0.146 | 0.07 | 0.055 | 0.101 |
| 43 | TM | Marrow | Batch | 0.004 | 0.151 | 0.059 | 0.153 | 0.779 | 0.334 | 0.414 | 0.414 |
| 44 | TM | Marrow | Cell type | 0.051 | 0.068 | 0.055 | 0.064 | 0.125 | 0.077 | 0.103 | 0.15 |
| 45 | TM | Muscle | Batch | 0.009 | 0.29 | 0.131 | 0.219 | 0.767 | 0.568 | 0.389 | 0.044 |
| 46 | TM | Muscle | Cell type | 0.068 | 0.076 | 0.078 | 0.071 | 0.105 | 0.09 | 0.083 | 0.099 |
| 47 | TM | Thymus | Batch | 0.008 | 0.175 | 0.52 | 0.539 | 0.748 | 0.71 | 0.635 | 0.023 |
| 48 | TM | Thymus | Cell type | 0.093 | 0.083 | 0.1 | 0.089 | 0.081 | 0.109 | 0.08 | 0.085 |
| 49 | TM | Tongue | Batch | 0.005 | 0.06 | 0.294 | 0.403 | 0.848 | 0.261 | 0.223 | 0.009 |
| 50 | TM | Tongue | Cell type | 0.164 | 0.167 | 0.164 | 0.164 | 0.153 | 0.165 | 0.154 | 0.176 |
| 51 | TM | Trachea | Batch | 0.009 | 0.153 | 0.104 | 0.2 | 0.661 | 0.463 | 0.233 | 0.119 |
| 52 | TM | Trachea | Cell type | 0.034 | 0.037 | 0.035 | 0.035 | 0.042 | 0.041 | 0.037 | 0.049 |
| 53 | Panc-reas | Pancreas_1 | Batch | 0.026 | 0.282 | 0.241 | 0.237 | 0.819 | 0.406 | 0.413 | 0.759 |
| 54 | Panc-reas | Pancreas_1 | Cell type | 0.039 | 0.038 | 0.039 | 0.039 | 0.05 | 0.04 | 0.037 | 0.065 |
| 55 | Panc-reas | Pancreas_2 | Batch | 0.008 | 0.133 | 0.143 | 0.189 | 0.829 | 0.257 | 0.321 | 0.733 |
| 56 | Panc-reas | Pancreas_2 | Cell type | 0.045 | 0.039 | 0.044 | 0.046 | 0.039 | 0.039 | 0.039 | 0.077 |
| 57 | Panc-reas | sub_Panc_reas_1 | Batch | 0.038 | 0.385 | 0.373 | 0.298 | 0.833 | 0.458 | 0.478 | 0.749 |
| 58 | Panc-reas | sub_Panc_reas_1 | Cell type | 0.041 | 0.039 | 0.045 | 0.049 | 0.086 | 0.039 | 0.047 | 0.083 |
| 59 | Panc-reas | sub_Panc_reas_2 | Batch | 0.114 | 0.584 | 0.511 | 0.486 | 0.755 | 0.629 | 0.679 | 0.756 |
| 60 | Panc-reas | sub_Panc_reas_2 | Cell type | 0.09 | 0.057 | 0.083 | 0.075 | 0.054 | 0.069 | 0.046 | 0.076 |

**Table S2:**
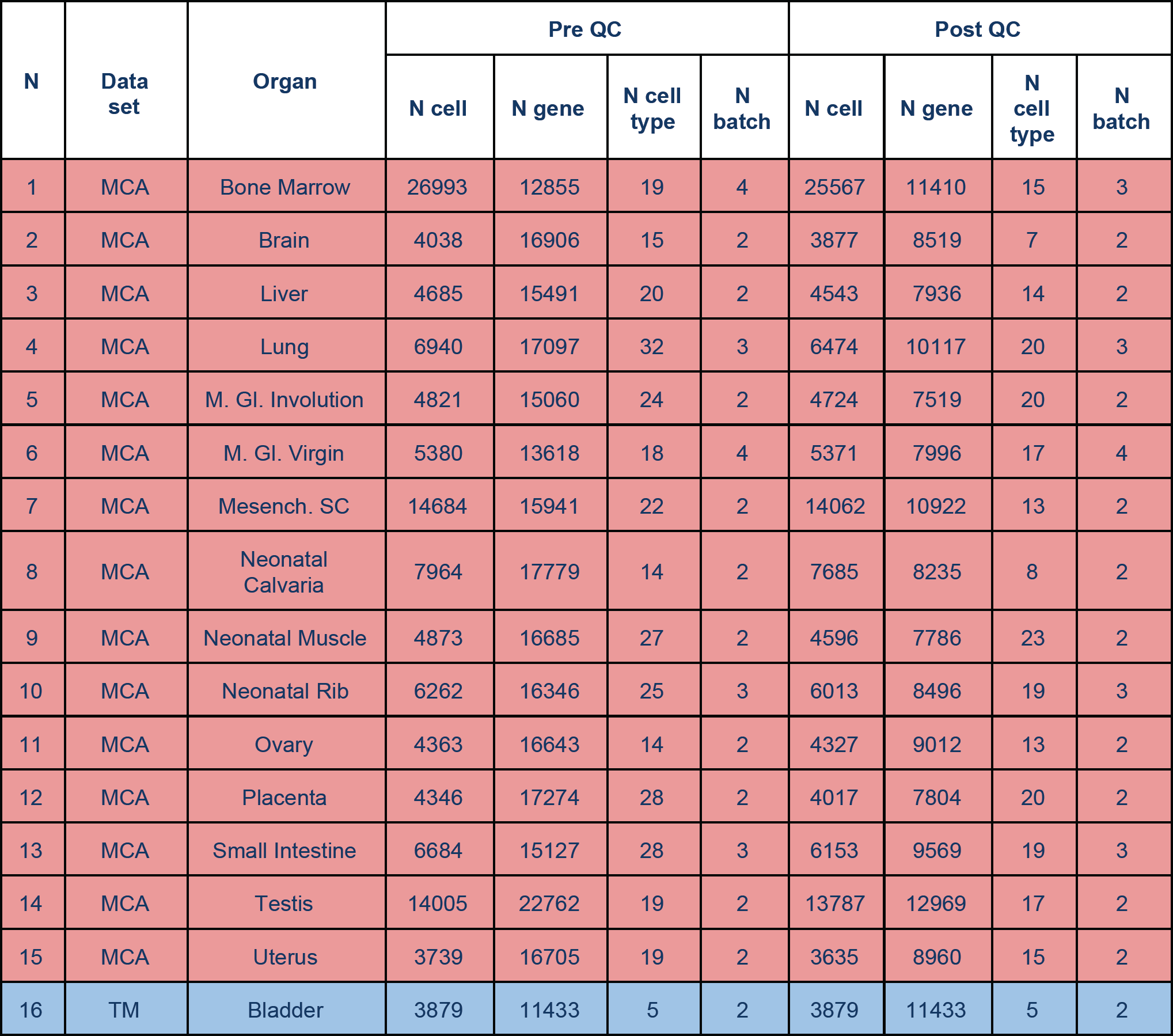
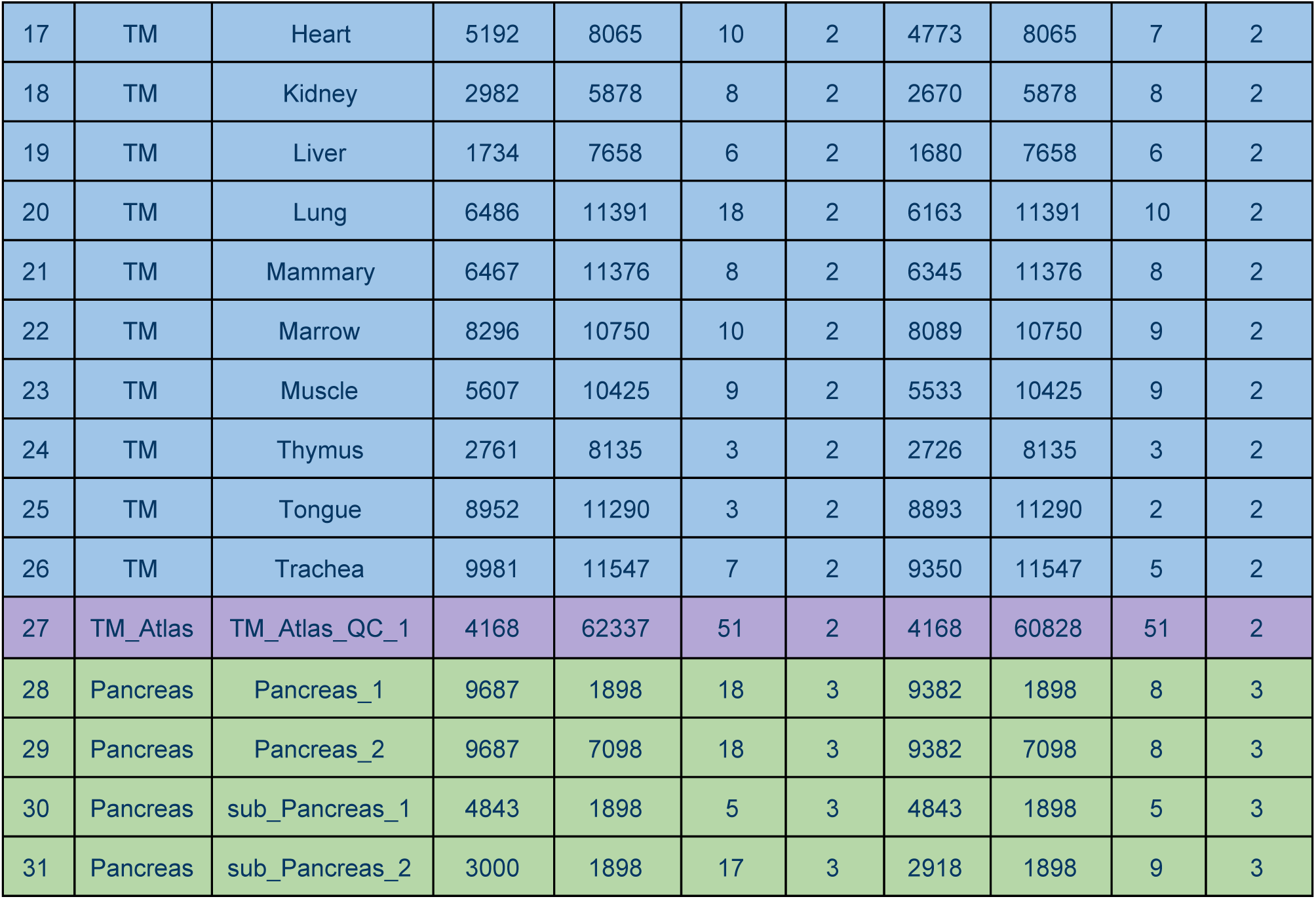
Summary statistics for the datasets considered in this study (MCA: Mouse Cell Atlas, TM: Tabula Muris).

| N | Data set | Organ | Pre QC |  |  |  | Post QC |  |  |  |
| --- | --- | --- | --- | --- | --- | --- | --- | --- | --- | --- |
|  |  |  | N cell | N gene | N cell type | N batch | N cell | N gene | N cell type | N batch |
| 1 | MCA | Bone Marrow | 26993 | 12855 | 19 | 4 | 25567 | 11410 | 15 | 3 |
| 2 | MCA | Brain | 4038 | 16906 | 15 | 2 | 3877 | 8519 | 7 | 2 |
| 3 | MCA | Liver | 4685 | 15491 | 20 | 2 | 4543 | 7936 | 14 | 2 |
| 4 | MCA | Lung | 6940 | 17097 | 32 | 3 | 6474 | 10117 | 20 | 3 |
| 5 | MCA | M. Gl. Involution | 4821 | 15060 | 24 | 2 | 4724 | 7519 | 20 | 2 |
| 6 | MCA | M. Gl. Virgin | 5380 | 13618 | 18 | 4 | 5371 | 7996 | 17 | 4 |
| 7 | MCA | Mesench. SC | 14684 | 15941 | 22 | 2 | 14062 | 10922 | 13 | 2 |
| 8 | MCA | Neonatal Calvaria | 7964 | 17779 | 14 | 2 | 7685 | 8235 | 8 | 2 |
| 9 | MCA | Neonatal Muscle | 4873 | 16685 | 27 | 2 | 4596 | 7786 | 23 | 2 |
| 10 | MCA | Neonatal Rib | 6262 | 16346 | 25 | 3 | 6013 | 8496 | 19 | 3 |
| 11 | MCA | Ovary | 4363 | 16643 | 14 | 2 | 4327 | 9012 | 13 | 2 |
| 12 | MCA | Placenta | 4346 | 17274 | 28 | 2 | 4017 | 7804 | 20 | 2 |
| 13 | MCA | Small Intestine | 6684 | 15127 | 28 | 3 | 6153 | 9569 | 19 | 3 |
| 14 | MCA | Testis | 14005 | 22762 | 19 | 2 | 13787 | 12969 | 17 | 2 |
| 15 | MCA | Uterus | 3739 | 16705 | 19 | 2 | 3635 | 8960 | 15 | 2 |
| 16 | TM | Bladder | 3879 | 11433 | 5 | 2 | 3879 | 11433 | 5 | 2 |
| 17 | TM | Heart | 5192 | 8065 | 10 | 2 | 4773 | 8065 | 7 | 2 |
| 18 | TM | Kidney | 2982 | 5878 | 8 | 2 | 2670 | 5878 | 8 | 2 |
| 19 | TM | Liver | 1734 | 7658 | 6 | 2 | 1680 | 7658 | 6 | 2 |
| 20 | TM | Lung | 6486 | 11391 | 18 | 2 | 6163 | 11391 | 10 | 2 |
| 21 | TM | Mammary | 6467 | 11376 | 8 | 2 | 6345 | 11376 | 8 | 2 |
| 22 | TM | Marrow | 8296 | 10750 | 10 | 2 | 8089 | 10750 | 9 | 2 |
| 23 | TM | Muscle | 5607 | 10425 | 9 | 2 | 5533 | 10425 | 9 | 2 |
| 24 | TM | Thymus | 2761 | 8135 | 3 | 2 | 2726 | 8135 | 3 | 2 |
| 25 | TM | Tongue | 8952 | 11290 | 3 | 2 | 8893 | 11290 | 2 | 2 |
| 26 | TM | Trachea | 9981 | 11547 | 7 | 2 | 9350 | 11547 | 5 | 2 |
| 27 | TM_Atlas | TM_Atlas_QC_1 | 4168 | 62337 | 51 | 2 | 4168 | 60828 | 51 | 2 |
| 28 | Pancreas | Pancreas_1 | 9687 | 1898 | 18 | 3 | 9382 | 1898 | 8 | 3 |
| 29 | Pancreas | Pancreas_2 | 9687 | 7098 | 18 | 3 | 9382 | 7098 | 8 | 3 |
| 30 | Pancreas | sub_Pancreas_1 | 4843 | 1898 | 5 | 3 | 4843 | 1898 | 5 | 3 |
| 31 | Pancreas | sub_Pancreas_2 | 3000 | 1898 | 17 | 3 | 2918 | 1898 | 9 | 3 |

